# Genetic fine-mapping from summary data using a non-local prior improves detection of multiple causal variants

**DOI:** 10.1101/2022.12.02.518898

**Authors:** Ville Karhunen, Ilkka Launonen, Marjo-Riitta Järvelin, Sylvain Sebert, Mikko J. Sillanpää

**Author notes:** **Contact info** Corresponding author: Ville Karhunen.

## Abstract

Genome-wide association studies (GWAS) have been successful in identifying genomic loci associated with complex traits. Genetic fine-mapping aims to detect independent causal variants from the GWAS-identified loci, adjusting for linkage disequilibrium (LD) patterns. The use of GWAS summary statistics and an adequate LD reference enable large sample sizes for fine-mapping, without direct access to individual-level data. We present *FiniMOM* (fine-mapping using a product inverse-moment priors), a novel Bayesian fine-mapping method for summarized genetic associations. For causal effects, the method uses a non-local inverse-moment prior, which is a natural prior distribution to model non-null effects in finite samples. A beta-binomial prior is set for the number of causal variants, with a parameterization that can be used to control for potential misspecifications in the LD reference. We test the performance of our method against a current state-of-the-art fine-mapping method SuSiE (sum-of-single-effects) across a range of simulated scenarios aimed to mimic a typical GWAS on circulating protein levels, and an applied example. The results show improved credible set coverage and power of the proposed method, especially in the case of multiple causal variants within a locus. The superior performance and the flexible parameterization to control for misspecified LD reference make *FiniMOM* a competitive alternative to other fine-mapping methods for summarized genetic data.

## 1 Introduction

Leveraging genetic associations from up to millions of individuals, genome-wide association studies (GWAS) have been widely used to find genomic loci associated with complex traits and disease liability [1]. Such genomic regions can be further prioritized to analyze disease aetiology and potential pharmacological targets in more detail [2].

Each genomic loci highlighted in a GWAS may harbor several causal variants (such as single-nucleotide polymorphisms [SNPs]), and the identification of these variants is important for a better understanding of the biological mechanisms underlying the trait of interest. The difficulty in dissecting the causal variants within a specific locus is compounded by the linkage disequilibrium (LD) patterns, as non-causal variants in LD with a true causal variant will also show associations with the trait of interest.

Fine-mapping methods aim to distinguish independent causal variants for a given trait within a specific genomic locus [3]. Assuming additive effects of individual variants, fine-mapping can be considered as a variable selection problem, where the aim is to identify the true causal signals from a candidate set of genetic variants [4], [5].

As the effect sizes of individual genetic variants are typically minuscule, large sample sizes are required in both GWAS and fine-mapping to obtain adequate statistical power to detect robust genetic associations. In addition to practical difficulties and privacy concerns in providing access to large-scale individual-level genetic data, summarized genetic associations from GWASs are increasingly publicly available. Therefore, the use of summary-level genomic data has become a standard in carrying out post-GWAS analyses [6]. Accordingly, many of the common fine-mapping methods are either compatible with, or developed for, GWAS summary statistics [7]–[16].

A key element in fine-mapping using summarized data is the correct specification of the LD structure underlying the locus-specific genetic associations. The most simple method of summary data fine-mapping via Bayes factors [7] makes a simplifying assumption of one causal variant per locus. This strategy has the benefit of not needing LD information, albeit the assumption itself may not be realistic [17]. However, if no in-sample LD information is available, care must be taken in how to obtain the LD reference. Ideally, the reference is derived from a sufficiently-sized sample [18] that is ancestrally similar to the population in the summary statistics, and with similar data quality control procedures applied [19]. In practice, this target may be very difficult to achieve [19].

Here, we propose a Bayesian fine-mapping method for quantitative traits based on non-local product inverse-moment (piMOM) priors using summary-level genetic associations. Originally proposed by Johnson & Rossell [20], non-local prior densities have zero density at the null parameter value, and the non-local product priors are independent products of such densities [21]. Such priors for regression coefficients have attractive properties of both theoretical and finite-sample performance for variable selection and prediction [21]–[23]. In the context of genomic analyses, non-local priors for binary, continuous, and time-to-event outcomes have been developed for individual-level data [24]–[26]. Prior to our work, no methods based on non-local priors have been applied on summary-level genetic data. Of note, analyses of summary-level genomic data provide an additional benefit of algorithms not scaling with the sample size.

The proposed method allows a flexible way to take the external LD information into account. We treat the model dimension as a parameter for which we assign a beta-binomial prior distribution. Such prior has been shown to provide optimal model selection in high-dimensional settings [27]. Crucially, our formulation of the prior allows the proposed method to be adjusted according to whether an in-sample or out-of-sample LD information is used.

We further apply a locally balanced proposal in our model selection posterior sampling algorithm, leading to optimal performance regarding the mixing time [28]. We demonstrate our method’s competitive performance to the current state-of-the-art fine-mapping method SuSiE (sum-of-single-effects regression) by different simulated scenarios and an applied example.

## 2 Methods

### 2.1 Statistical model

Let ***Y*** = (*y*_1_ … *y*_*N*_)^*T*^ be a vector of the mean-centered observed values for a quantitative trait of interest, and ***X*** a standardised (mean = 0; standard deviation = 1) genotype matrix of size *N* × *P* for genetic variants within a specific genomic locus. Given a linear model

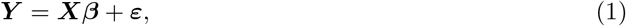

where 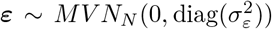, our aim is to identify the causal variants *x*_*j*_ for which *β*_*j*_ ≠ 0. In other words, we aim to identify a model (or a *causal configuration*) ***m***, which only includes the *k* causal variants, from a set of candidate models ℳ of maximum size (dimension) *K*. Typically, the magnitude of variants included in a fine-mapping application vary from *P* ≈ 10^2^ to *P* ≈ 10^3^, and it is also reasonable to assume that the true model dimension *k* ≤ *K* ≪ *P*.

In the absence of individual-level genetic data, we resort to the use of GWAS summary statistics, where the outcome is regressed on each genetic variant separately, with population stratification adequately taken into account, and with further adjustments for additional covariates, such as sex, age or technical covariates, carried out where appropriate [29]. We require that the coefficient estimates 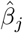 and their standard errors 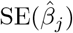 are available in the GWAS summary statistics. Furthermore, we assume that there is a good estimate available for the LD matrix ***R***_*P* ×*P*_, and that the effect allele frequencies (EAFs) for each variant and Var(*Y*) are known.

Assuming a large *N* and small *β*_*j*_ (both of which are usually the case in GWAS), we follow the proposal by Zhu & Stephens [30] and write the likelihood for the observed effect estimates 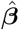 as

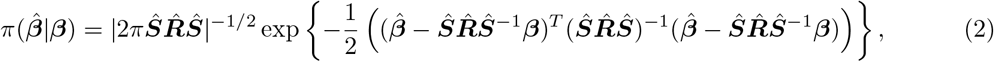

where 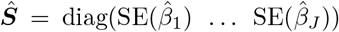 and 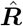 is an estimate of LD matrix ***R***. If the summary statistics are given in per-allele units, these can be trivially transformed into standardized units by assuming a Hardy-Weinberg equilibrium for the variants and multiplying both 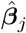 and 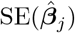 by 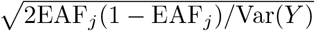.

#### 2.1.1 Non-local prior for effect size *β*

A Gaussian density is commonly used for an effect size prior of a causal variant. Such zero-centered, i.e. ‘local’, priors have the maximum density for the causal effect size at the null parameter value. However, when conditioning on model ***m*** which only includes the causal variant(s), such prior densities may seem counter-intuitive due to unnecessary strong shrinkage of the causal variants towards the null value. Therefore, conditioned on model ***m***, we set a non-local product inverse-moment (piMOM) prior for the causal effect size vector ***β***:

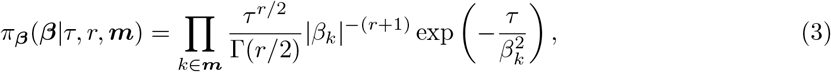

for *τ >* 0 and *r* = 1, 2, This prior is a product of independent inverse moment prior densities for each component of the parameter vector in the model. As per the definition of a non-local prior, the density value at zero is 0, which is a natural characteristic for the effect size prior. The product formulation of the prior ensures that the density is zero if any of the components in the parameter vector is zero, which leads to a strong penalty on the parameters [21]. Of note, in equation (3), we also implicitly condition on the model dimension *d*.

Parameter *r* controls the tail behavior of the distribution. Selecting *r* = 1 leads to Cauchy-like tails, which are known to protect against the potential over-shrinkage of large effect sizes for sparse signals [31].

The parameter *τ* controls the spread of the effects away from 0, and it can be set so that with a selected probability, the effect sizes are above a specific threshold. For example, with a prior belief that ℙ (|*β*_*j*_| *>* 0.05) = 0.99, we get *τ* = 0.0083 [25].

Figure 1 depicts a comparison of a marginal inverse moment density with a Gaussian density, which is commonly used as an effect size prior in genetics [32]. The inverse moment prior possesses heavier tails which allow for less shrinkage of large effect sizes, and while its density vanishes as *β*_*j*_ → 0, there is still a non-zero prior probability to detect small but non-zero effect sizes.

**Figure 1:**
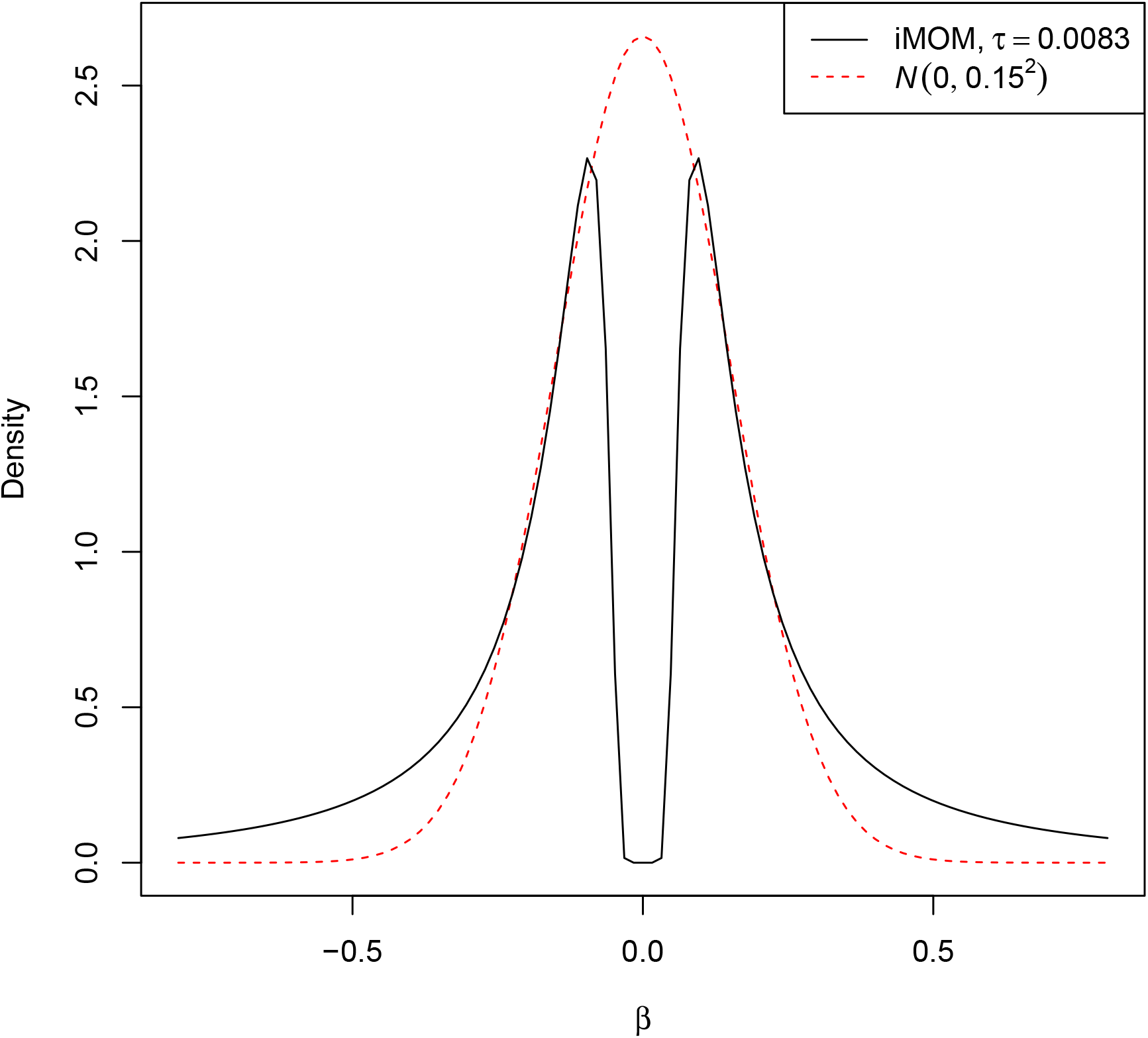
Comparison of a marginal inverse moment (iMOM) distribution with *τ* = 0.0083 and a Gaussian (*N*) distribution with mean 0 and variance 0.15^2^.

#### 2.1.2 Prior for model dimension and accounting for potential LD misspecifications

As a prior for model dimension *d* (i.e., for the number of causal variants), we assign a Beta-binomial distribution:

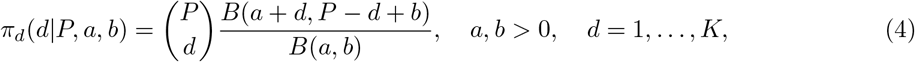

where *B*(·) is the Beta function. This prior arises from a binomial distribution for the model dimension with parameters *P* and *q*, where *q* is being assigned a Beta(*a, b*) distribution. As fine-mapping is typically applied in situations with at least one causal variant, we set ℙ(*d* = 0) = 0. The model dimension is also restricted by a maximum model dimension *K* ≪ *P*, with ℙ(*d* = *l*) = 0 for *l* = *K* + 1, …, *P*. The beta-binomial prior is consistent with the family of priors suggested by Castillo et al. [27] for recovering the correct sparse model in high-dimensional regression.

One choice for the parameters *a* and *b*, as discussed in Castillo et al. [27] and Castillo & van der Vaart [33], is to set *a* = 1 and *b* = *P* ^*u*^, *u >* 1. The hyperparameter *u* controls the concentration of the probability mass of the model dimension prior, in that larger *u* prioritizes smaller models *a priori* (Supplementary Figure 1). Earlier work on summary data fine-mapping has shown that a misspecified LD matrix is likely to lead to more false positives [18]. Therefore, a natural and an adaptive way to take into account the potential misspecification of the LD matrix is by increasing *u*, which places more prior probability mass to models of smaller dimension. This protects against false positives in the posterior while allowing the data to pick up strong signals of multiple causal variants, provided that the data strongly favor such models. Importantly, in the case of in-sample (or otherwise highly accurate) LD matrix being available, such strong shrinkage towards sparse models is not necessarily needed, and therefore *u* can be set to a smaller value, thus not compromising power at the expense of the false positives.

#### 2.1.3 Posterior inference

Let *π*_***m***_(***D***) denote the the marginal likelihood of the data under model ***m***, again implicitly conditioning on the model dimension *d*. To obtain the posterior probability for model ***m*** of dimension *d*, given data 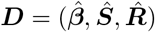, we can apply Bayes’ rule and obtain

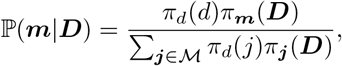

where *j* refers to the dimension of model ***j***. When comparing different models, the denominator is the same for all models and can be cancelled out in the algebra. Similarly as in Johnson & Rossell [21] and Nikooienejad et al. [24], *π*_***m***_(***D***) can be computed via Laplace’s method:

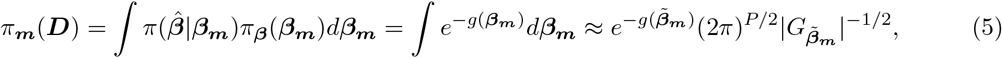

where ***β***_***m***_ refers to the parameter vector corresponding to the model ***m***,

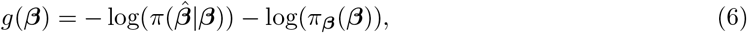

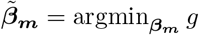 and 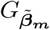 is the Hessian of *g* evaluated at 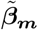.

Plugging in equations (2) and (3) into (6) and dropping the constant terms with respect to the parameters, we have

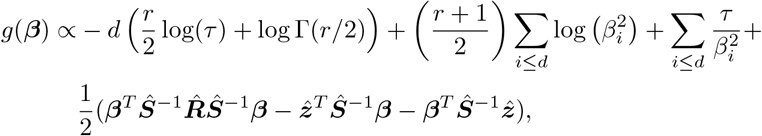

where 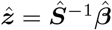, and the Hessian

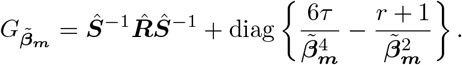

The value of 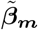 can be obtained by a numerical optimization method. Of note, computing the marginal likelihood does not involve calculating the inverse of 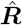. Therefore, this method is flexible even in the case of non-invertible LD matrices (see also Zou et al. [16]), as well as remaining computationally efficient.

#### 2.1.4 Markov chain Monte Carlo sampling scheme

To generate dependent samples from the posterior distribution, we propose the following Markov chain Monte Carlo (MCMC) sampling scheme:

1. Choose initial model ***m***^curr^.
2. For *i* = 1, …, *n*_iter_
  a. To create a proposal model ***m***^prop^, randomly select to either add, delete, or swap one of the active variables in ***m***^curr^:
    i. Add variable with probability *p*_add_ that is proportional to the squared residuals of the current model: *p*_add_ ∝ |*X*^*T*^ *ε*|^2^, where 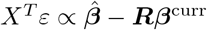
    ii. Delete variable with uniform probability from all active variables.
    iii. Swap: select a variable to be swapped to inactive with uniform probability from all active variables in ***m***^curr^ – select the variable to be swapped to active with probability *p*_swap_ that is proportional to the squared correlation with the variable to be swapped to inactive: 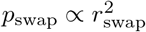
  b. Using the proposed model ***m***^prop^, compute acceptance probability

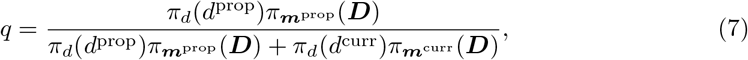

where *d*^prop^ and *d*^curr^ are the dimensions of models ***m***^prop^ and ***m***^curr^, respectively.
  c. Sample *u* ∼ *U* (0, 1), and if *q > u*, then set ***m***^curr^ = ***m***^prop^.

In the ‘add’-step, we give the largest probabilities to variables that have the largest correlations with the residuals of the current model. This bears similarities to the screening method proposed in Shin et al. [23]. Our method allows a non-zero probability for all variables, instead of considering only the variables that correlate highly with the residuals of the current model.

In the ‘swap’-step, we prioritize variants with high correlations with a variant already existing in the model. This prevents the model from deviating far from an already good fit, in that the variant added to the model is likely in high LD with a variant to be replaced. This also provides a natural and efficient way to take into account the uncertainty of a true causal variant among a group of variants in very high LD (see also Section 2.1.6 for further considerations of extremely high LD).

The acceptance probability in equation (7) is also called the Barker proposal [34], suggested for non-local priors in Johnson & Rossell [21]. Furthermore, Zanella [28] showed this to be the optimal proposal regarding the mixing time in the case of binary indicators. Accordingly, by examining the trace plots for the realizations of the model dimension parameter in the simulations (Section 2.2), we found similar convergence for Markov chains of length 12,500 with a 2500 burn-in (i.e. 10,000 samples from the posterior) as for Markov chains of length 60,000 with a 10,000 burn-in (50,000 samples).

In equation (7), the model dimension prior *π*_*d*_ and the marginal likelihood *π*_***m***_ refer to those in equations (4) and (5), respectively. The interest is in whether the variants are causal or not, or in other words, whether they are included in model ***m***. Therefore, we have integrated out all other parameters (that is, the effect sizes *β*) in equation (5) using Laplace’s method. The proposed sampling scheme is related to reversible jump MCMC algorithm [35], however our formulation and use of Laplace’s method avoids complicated sampling from varying-dimensional model space.

The proposed fine-mapping method *FiniMOM* (fine-mapping using a product inverse-moment prior) with the described sampling scheme is implemented in a freely available R package: https://github.com/vkarhune/finimom.

#### 2.1.5 Credible sets

We adopt the proposed approach in Lee et al. [36] and Wang et al. [15] and treat credible sets as the main tool for posterior inference. A credible set at level *α* ∈ (0, 1) is defined as a set of variants that contain a true causal variant with probability larger than *α*.

Via credible sets, we can present the uncertainty in both (i) the number of causal variants that are supported by the data, and (ii) the identification of a true causal variant from a candidate set of variants for a specific set. We can directly use the posterior distribution of model dimension, ℙ(*d* = *l*|***D***), 1 ≤ *l* ≤ *K*, to obtain the posterior probabilities for the supported number of signals *l*, and equivalently, the number of credible sets. The credible sets are then created at coverage *α*_*l*_, which represent the coverage of *α* conditioned on *l* signals. In addition to the credible sets, posterior inclusion probabilities (PIPs) for each variant PIP_*j*_ = ℙ(*β*_*j*_ ≠ 0|***D***), *j* = 1, …, *J*, can be calculated as the proportion of the posterior samples where variant *j* is included in the model.

#### 2.1.6 Clumping extremely highly correlated variants

Extremely highly correlated variants, such as those with pairwise *r*^2^ *>* 0.99, cannot be easily distinguished by any fine-mapping method. Therefore, we have added an option to clump such variants to be treated as one ‘cluster’ in the posterior sampling, with the lead variant (i.e. the one with the lowest marginal *p*-value) representing this group of variants in the sampling, yielding a speed improvement as a by-product. Consequently, any credible set that would include the lead variant from a cluster of clumped variants would also include all other variants from the corresponding cluster.

#### 2.1.7 Consistency check for out-of-sample LD reference

When using out-of-sample LD reference, there is a possibility of inconsistencies in that the observed test statistic does not correspond to the estimated LD between the variants; Zou et al. [16] give a toy example of such scenario of two variants, with 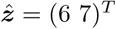 and 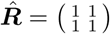.

We present an option to distinguish such variants. Considering a cluster of variants in extremely high LD, we assume the variant *C* with the lowest p-value as the true causal variant. As per [9], we assume that the observed test statistic for any non-causal variant *N* in this cluster depends solely on its correlation with *C*. The joint distribution of the marginal *z* scores for *C* and *N* is therefore a multivariate Gaussian distribution:

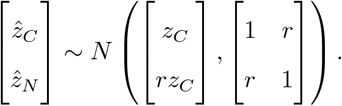

For testing equality of the observed and expected z-scores for *N*, we obtain a test statistic

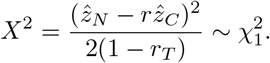

In the denominator, we approximate *r* with *r*_*T*_, the threshold used to define extremely highly correlated variants. If the observed 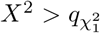, the LD structure is not considered to approximate the *z* scores, and the variants are assigned to separate clusters. By default, we use 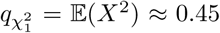. This provides additional robustness to the clumping and serves as a check for LD misspecification of tightly-linked variants. We highlight that none of the variants are excluded from fine-mapping, but simply treated as separate, non-clumped clusters in the sampling phase. In the toy example, our method would distinguish the misspecified LD structure and treat the variants separately. Similar to SuSiE [16], our method also strongly prefers the second variant to be the causal one.

### 2.2 Simulation study

We conducted a simulation study to investigate the accuracy of our proposed method. The genotype data used for simulations were obtained from two Finnish population-based pregnancy-birth cohorts, Northern Finland Birth Cohort 1966 (NFBC1966) [37]–[39] and Northern Finland Birth Cohort 1986 (NFBC1986) [40], [41].

Genotyping and quality control for both datasets are detailed elsewhere [42], [43]. Briefly, genotyping was done using Illumina HumanCNV-370DUO Analysis BeadChip (Illumina, California, USA) for NFBC1966 and Illumina HumanOmniExpressExome-8v1.2 platform for NFBC1986. After sample quality control procedures and consent withdrawals, genomic data were available for 5400 and 3743 individuals in NFBC1966 and NFBC1986, respectively. Both datasets were Imputed to Haplotype Reference Consortium panel, and the autosomal, biallelic markers were filtered for minor allele frequency 0.01, imputation quality *R*^2^ *>* 0.5 and Hardy-Weinberg equilibrium *p*-value *>* 10^−12^.

In the simulation studies, the genotype data from NFBC1966 were used for phenotype simulations to create the summary-level data, and genotype data from NFBC1986 for estimating the out-of-sample LD matrix. These data come from the same geographical region with no overlapping individuals, therefore the different datasets provide ideal out-of-sample LD references for each other. Due to the larger sample size in NFBC1966, we used this study to simulate the phenotype and the summary data, and NFBC1986 for out-of-sample LD reference.

We randomly selected five protein-coding genes of varying size from different chromosomes, and selected the region *±* 100kb around each gene for fine-mapping. The variants were filtered for availability in both NFBC1966 and NFBC1986 datasets, resulting in the number of variants in the fine-mapped loci varying from 387 to 2996 (Supplementary Table 1 and Supplementary Figures 2–6).

**Table 1:** Estimates and 95% confidence intervals (CI) for mean credible set coverage, mean credible set power, and median credible set size across all simulated scenarios. The CIs are based on binomial distribution for coverage and power, and 1000 bootstrap samples for median size. FiniMOM: fine-mapping using inverse-moment prior; SuSiE: sum of single effects; LD: linkage disequilibrium.

| Method | LD matrix | Coverage | Power | Median size |
| --- | --- | --- | --- | --- |
| FiniMOM | In-sample | 0.912 (0.904 – 0.920) | 0.542 (0.531 – 0.553) | 12 (12 – 13) |
| FiniMOM | Out-of-sample | 0.906 (0.897 – 0.915) | 0.507 (0.496 – 0.518) | 11 (11 – 12) |
| SuSiE | In-sample | 0.890 (0.881 – 0.899) | 0.492 (0.481 – 0.503) | 11 (10 – 11) |
| SuSiE | Out-of-sample | 0.887 (0.877 – 0.896) | 0.492 (0.481 – 0.503) | 11 (10 – 12) |

Using NFBC1966 genotype data, we simulated a quantitative trait under the model in equation (1) with *C* causal variants explaining a total of *ϕ* of variance in the phenotype, with *C* ∈ {1, 2, 5} and *ϕ* ∈ {0.015, 0.03} as follows:

1. Select indices for the causal variants *C* uniformly from {1, …, *P*}, with the constraint that the maximum correlation between the causal variants is 0.95.
2. Draw the effect sizes with the constraint that the power to detect all individual causal effect sizes was *>* 0.8 at significance level = 0.05:
  a. Draw *C* effect sizes *b*_*c*_ from *N* (0, 1).
  b. Scale the *b*_*C*_ to have the desired total variance explained by simulating an unscaled phenotype *Y* ^*′*^ with

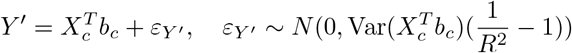

and then rescaling

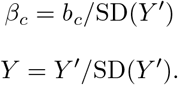
  c. Return to (a) if any of *β*_*c*_ do not achieve the desired power.

The parameters used in the simulations represent plausible scenarios for protein quantitative trait loci (pQTL) analyses [44]. The simulated phenotype was regressed separately on all variants (standardized to mean = 0 and variance = 1) within the locus assuming an additive model. The resulting summary statistics (i.e. association estimates and their standard errors) and an LD matrix (see below) were given as inputs to the considered fine-mapping methods.

To investigate the influence of the choice of the LD matrix 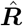, we compared the performance of using in-sample LD matrix (i.e. calculated using NFBC1966 genotype data) and out-of-sample LD matrix, calculated using NFBC1986 genotype data. We clumped the variants at *r*^2^ = 0.99, and applied the LD inconsistency check when out-of-sample LD matrix was used.

Each scenario was repeated 100 times. For all simulations, *τ* = 0.0083, *r* = 1, and the maximum number of causal variants *K* = 10. We varied the hyperparameter *u* ∈ {1.05, 1.25, 1.50, 1.75} to examine the impact of the model dimension prior on the results. MCMC was run for 12,500 iterations, of which we excluded the first 2500 as a “burn-in”, and used the last 10,000 for posterior inference.

The two main performance measures were the credible set coverage (the proportion of the credible sets containing a true causal variant) and power (the proportion of true causal variants included in a credible set). We also evaluated the median size of the credible sets, the sets being ideally as small as possible with adequate coverage and power. The sampling variability of the estimates in the replication results is quantified by 95% confidence intervals, calculated using binomial distribution for coverage and power, and by 1000 bootstrap samples for median model dimension.

We compare our method with the current state-of-the-art method of sum-of-single-effects (SuSiE) regression using summary statistics, which has been shown to perform well against many other fine-mapping methods [15], [16]. The SuSiE method applies a ‘sum-of-single-effects’ prior:

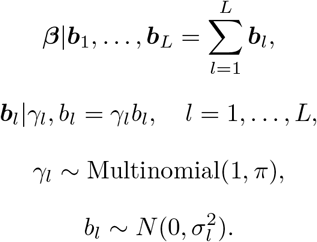

The overall vector of effect sizes ***β*** is a sum of *L* effect size vectors with exactly one non-zero element, with a Gaussian prior for the non-zero effect size. The model fitting is done via Iterative Bayesian Stepwise Selection (IBSS) described in Wang et al. [15], and Zou et al. [16] extended its adaptation for summarized data. We applied SuSiE with *L* = 10 and with the effect size prior variance 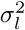 estimated from the data.

### 2.3 Applied real data example

We illustrate the performance of our method in real data for genetic associations of interleukin-18 (IL18), a pro-inflammatory cytokine that stimulates several cell types as an inflammatory factor [45]. The source GWAS summary statistics on IL18 are based on the analysis of 3,675 individuals in three Finnish cohorts [46], [47], which identified three loci with at least one variant associated with circulating IL18 levels at *p <* 5 × 10^−8^. These three loci (*±* 1Mb from the variant with the lowest *p*-value) were selected for fine-mapping, in which we applied FiniMOM with *τ* = 0.0083, *r* = 1, *u* = 1.75 and *K* = 10, and SuSiE (version 0.11.42) with *L* = 10 and 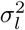 estimated from the data. NFBC1966 genotype data was used to generate the (out-of-sample) LD matrix. We compare the variants contained in the credible sets given by each method, and further assess the marginal associations of these variants in an external GWAS on IL18, conducted in up to 19,195 individuals of European ancestries [48].

The analyses were conducted using R software. The scripts for simulations are available at: https://github.com/vkarhune/finimomSimulations.

## 3 Results

### 3.1 Analysis of simulation replicates

The FiniMOM simulation results across varying values of hyperparameter *u* are presented in Figures 2 and 3. The credible set coverage increased with larger values of *u*, with *u* = 1.05 being the weakest in terms of credible set coverage across different numbers of causal variants, the variance explained by the causal variants, and whether in-sample or out-of-sample LD matrix was used. In general, the results were similar for other tested values of *u*. As expected, using the in-sample LD consistently outperformed the use of an LD matrix from a reference panel, however this difference diminished with increasing values of *u*.

**Figure 2:**
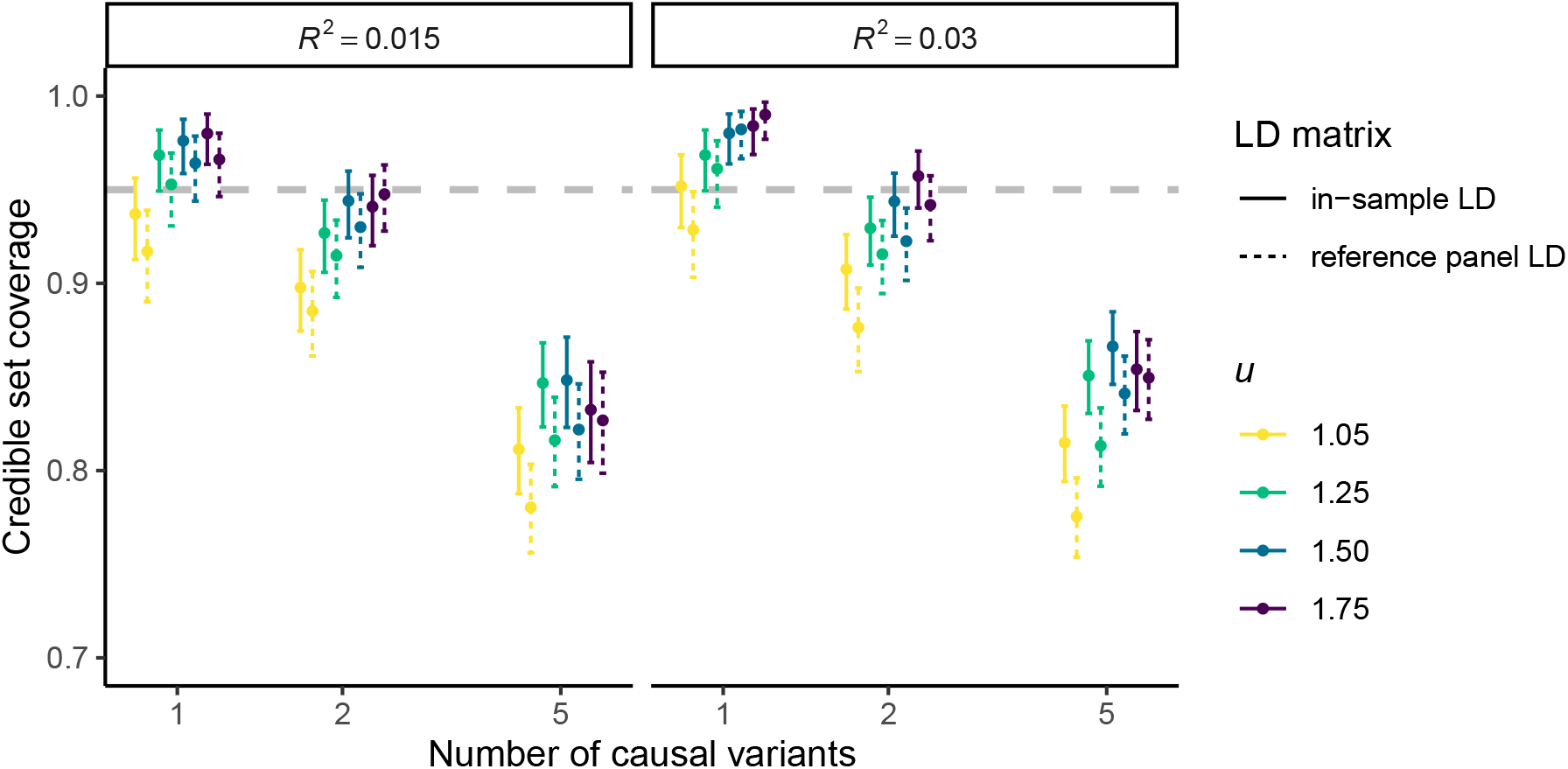
95% credible set coverage and their 95% confidence intervals in the simulation study (calculated over 100 simulation replicates) using FiniMOM with different values for hyperparameter *u*, which controls the prior for model dimension. Larger values of *u* refer to stronger priors toward smaller dimensions. The grey dashed line represents the nominal 95% target coverage. LD: linkage disequilibrium.

**Figure 3:**
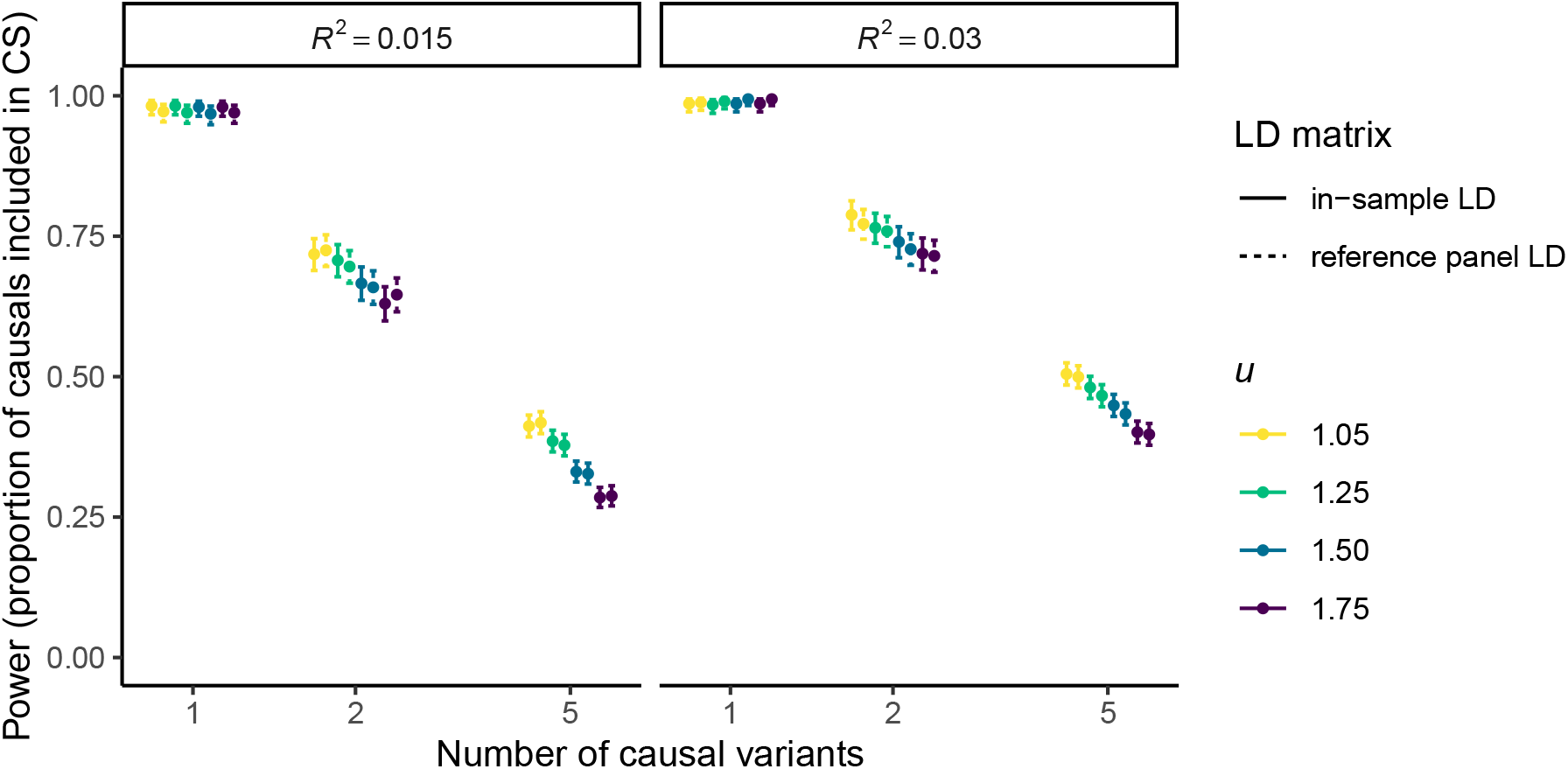
Credible set power and their 95% confidence intervals in the simulation study (calculated over 100 simulation replicates) using FiniMOM with different values for hyperparameter *u*, which controls the prior for model dimension. Larger values of *u* refer to stronger priors toward smaller dimensions. CS: credible set; LD: linkage disequilibrium.

All values of *u* had excellent power in the scenarios of one causal variant (Figure 3). When there were multiple causal variants, we saw the opposite effect compared to the coverage results, that is, the power was higher for smaller values of *u*. This is explained by smaller *u* allowing more prior probability mass for larger models, which leads to FiniMOM being more sensitive in finding solutions with larger number of causal variants. This comes at the expense of false positive credible sets, consequently decreasing the coverage.

Based on the investigation of FiniMOM performance using different values of *u*, we carried out the comparison with SuSiE fine-mapping using *u* = 1.50 for in-sample LD and *u* = 1.75 for out-of-sample LD. These values were selected as a compromise of the ideal credible set coverage and power.

The comparison of credible set coverage between FiniMOM and SuSiE is shown in Figure 4. While the differences within each scenario were mostly minor, FiniMOM produced consistently larger credible set coverage than SuSiE. The most pronounced differences were seen in the cases with five causal variants. Interestingly, FiniMOM with out-of-sample LD was no worse than SuSiE with in-sample LD in any of the scenarios.

**Figure 4:**
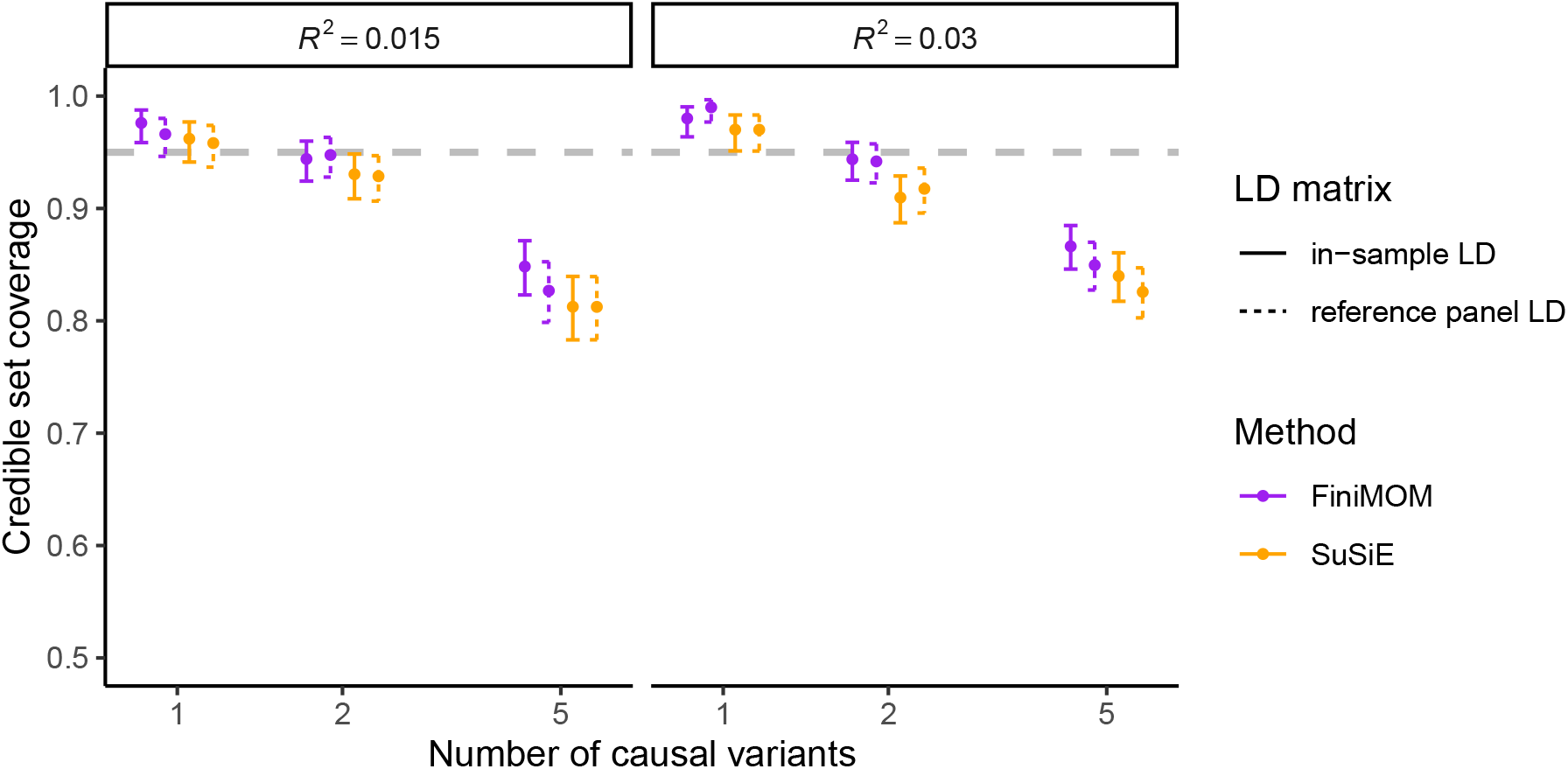
Comparison of credible set coverage and their 95% confidence intervals for FiniMOM and SuSiE in the simulation study (calculated over 100 simulation replicates). The grey dashed line represents the nominal 95% target coverage. LD: linkage disequilibrium; FiniMOM: Fine-mapping using inverse-moment priors; SuSiE: sum of single effects.

FiniMOM also showed better statistical power to detect multiple causal variants than SuSiE (Figure 5). As was the case for coverage, the largest differences in favour of FiniMOM were detected in the five-causal-variant scenarios. The median credible set sizes were similar or marginally larger in FiniMOM (Figure 6).

**Figure 5:**
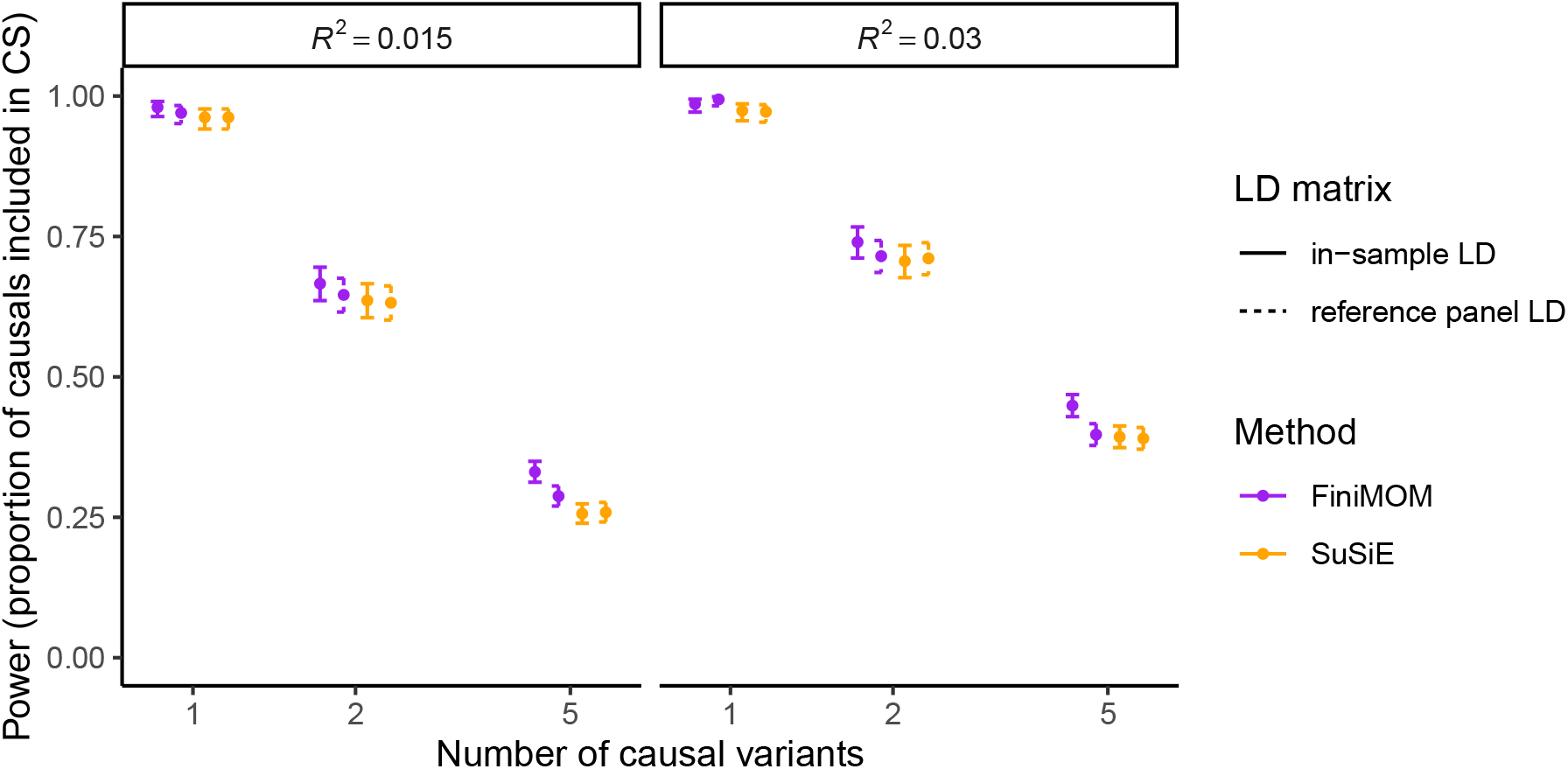
Comparison of credible set power and their 95% confidence intervals for FiniMOM and SuSiE in the simulation study (calculated over 100 simulation replicates). LD: linkage disequilibrium; FiniMOM: Fine-mapping using inverse-moment priors; SuSiE: sum of single effects.

**Figure 6:**
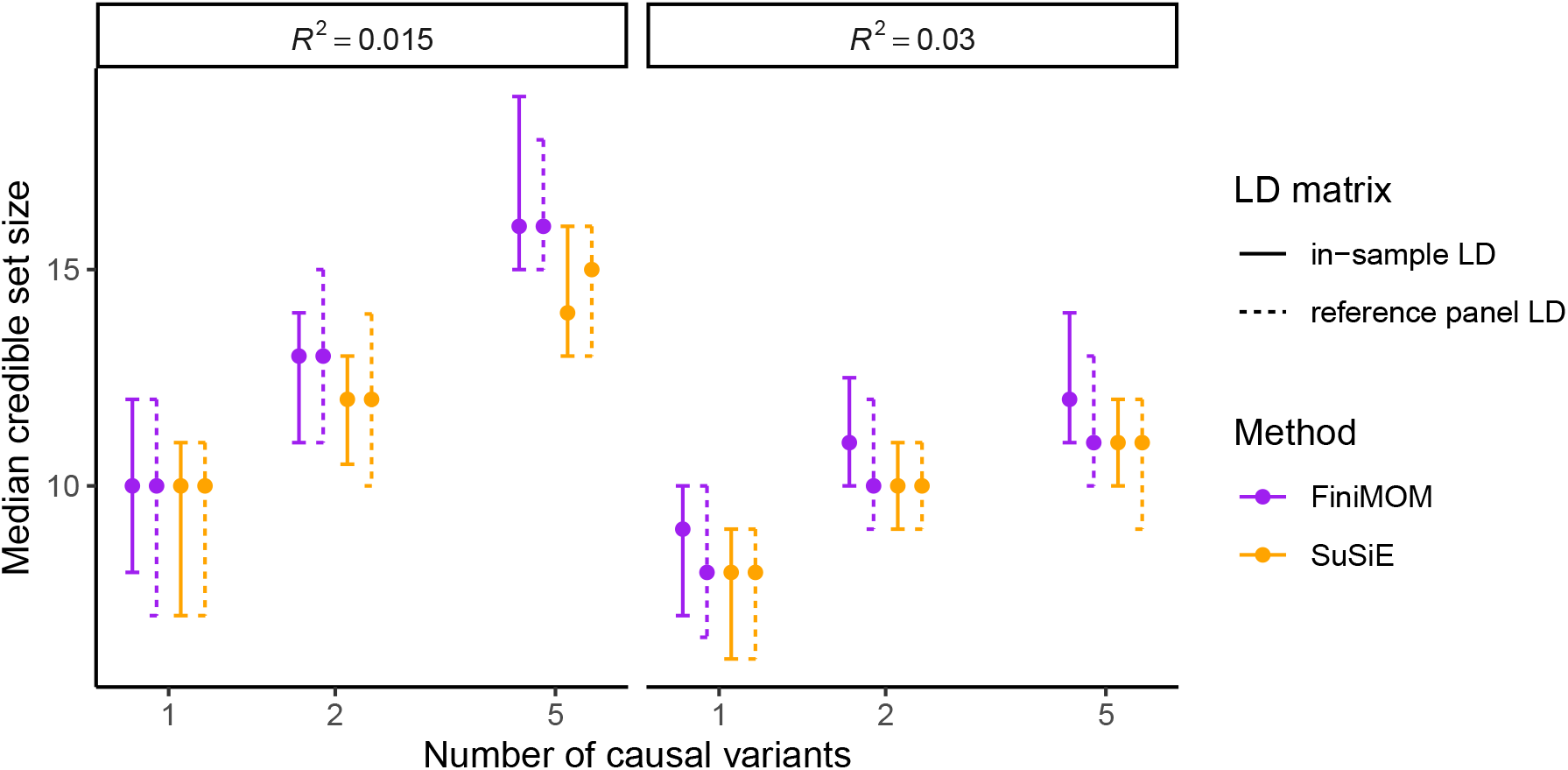
Comparison of credible set size (median and 95% confidence intervals) for FiniMOM and SuSiE in the simulation study (calculated over 100 simulation replicates). LD: linkage disequilibrium; FiniMOM: Fine-mapping using inverse-moment priors; SuSiE: sum of single effects.

The coverage and power estimates across all simulation scenarios (different number of causal variants and different values of *R*^2^) are given in Table 1. FiniMOM had better coverage and power for both in-sample LD and out-of-sample LD than SuSiE, with similar or marginally larger median credible set size.

### 3.2 Applied example

We then ran an applied example using GWAS summary statistics on circulating IL18 levels [47]. We chose three genomic regions – *NLRC4, RAD17* and *BCO2* – harbouring variants at *p <* 5 × 10^−8^ (*±* 1Mb window from the lead variant) for fine-mapping (Table 2), and investigated the variants in the produced credible sets in a separate GWAS on IL18 levels [48] for replication (“replication GWAS”). The run times (using Intel Xeon processor running at 2.1 GHz) for *NLRC4, RAD17* and *BCO2* loci were 81s, 66s and 60s, respectively. For SuSiE, the same run times were 7.8s, 3.6s and 6.2s, respectively.

**Table 2:** Genomic regions used in the applied example.

| Lead variant | Nearest gene | Genomic region (hg19) | Number of variants |
| --- | --- | --- | --- |
| rs385076 | *NLRC4* | chr2:31,489,851–33,489,851 | 6511 |
| rs17229943 | *RAD17* | chr5:67,682,536–69,682,536 | 3043 |
| rs12420140 | *BCO2* | chr11:111,071,294–113,071,294 | 5862 |

The results for IL18 are summarized in Figures 7, 8 and 9. Both methods selected the same number of credible sets for each loci, with minor deviations in the numbers of causal variants. The uncertainty in the number of credible sets is provided only by FiniMOM method (Table 3).

**Table 3:**
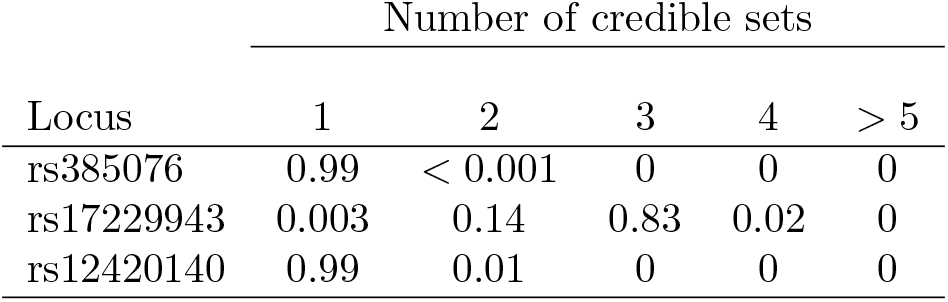
FiniMOM posterior distribution for the number of credible sets in each locus.

**Figure 7:**
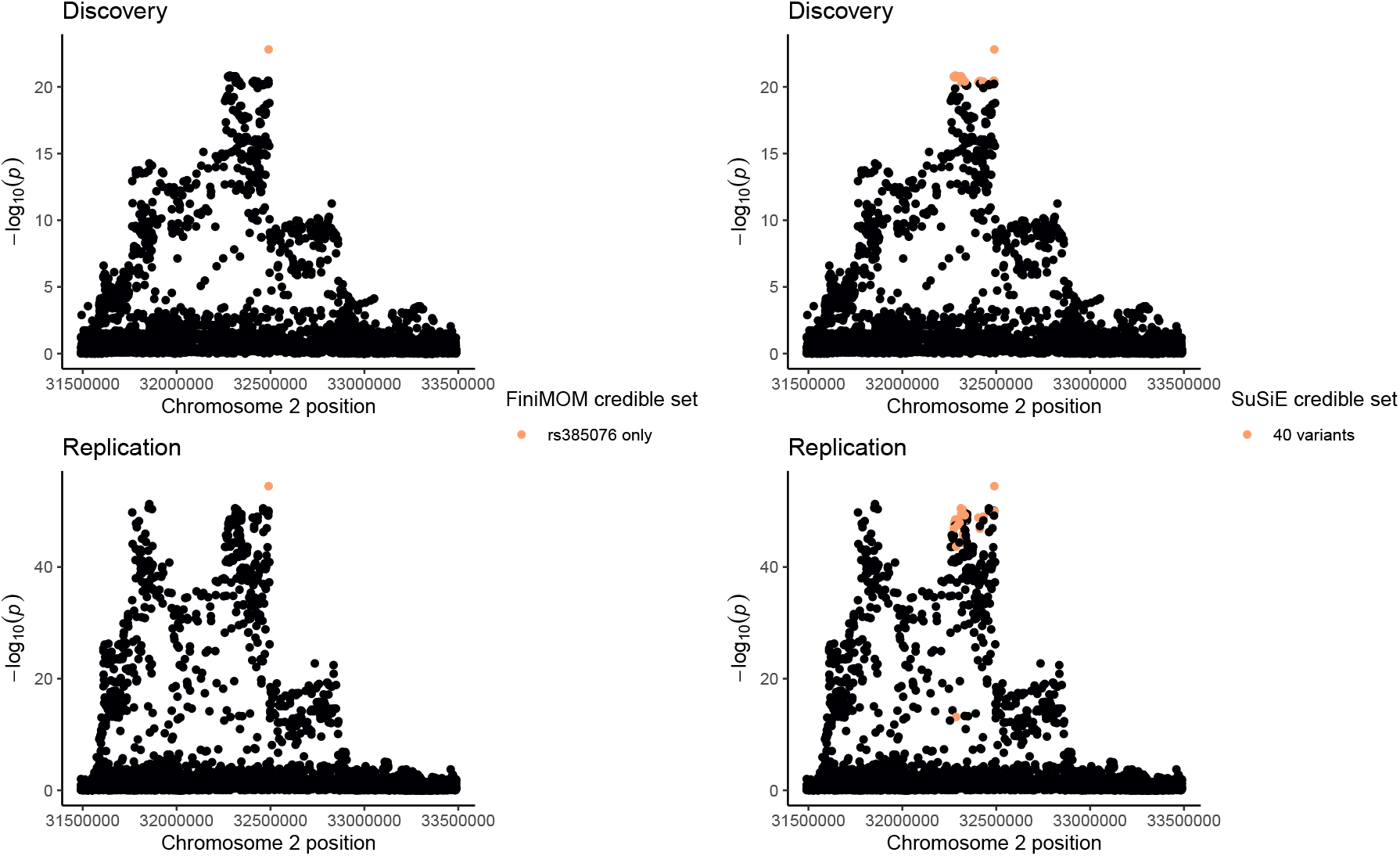
Locus plot of genetic associations (− log_10_(*p*)) per each variant within *NLRC4* locus (± 1Mb from rs385076 variant), with credible sets highlighted for FiniMOM (left panels) and SuSiE (right panels) in the discovery GWAS (top panels) and in the replication GWAS (bottom panels). FiniMOM: fine-mapping using inverse-moment prior; SuSiE: sum of single effects; GWAS: genome-wide association study.

**Figure 8:**
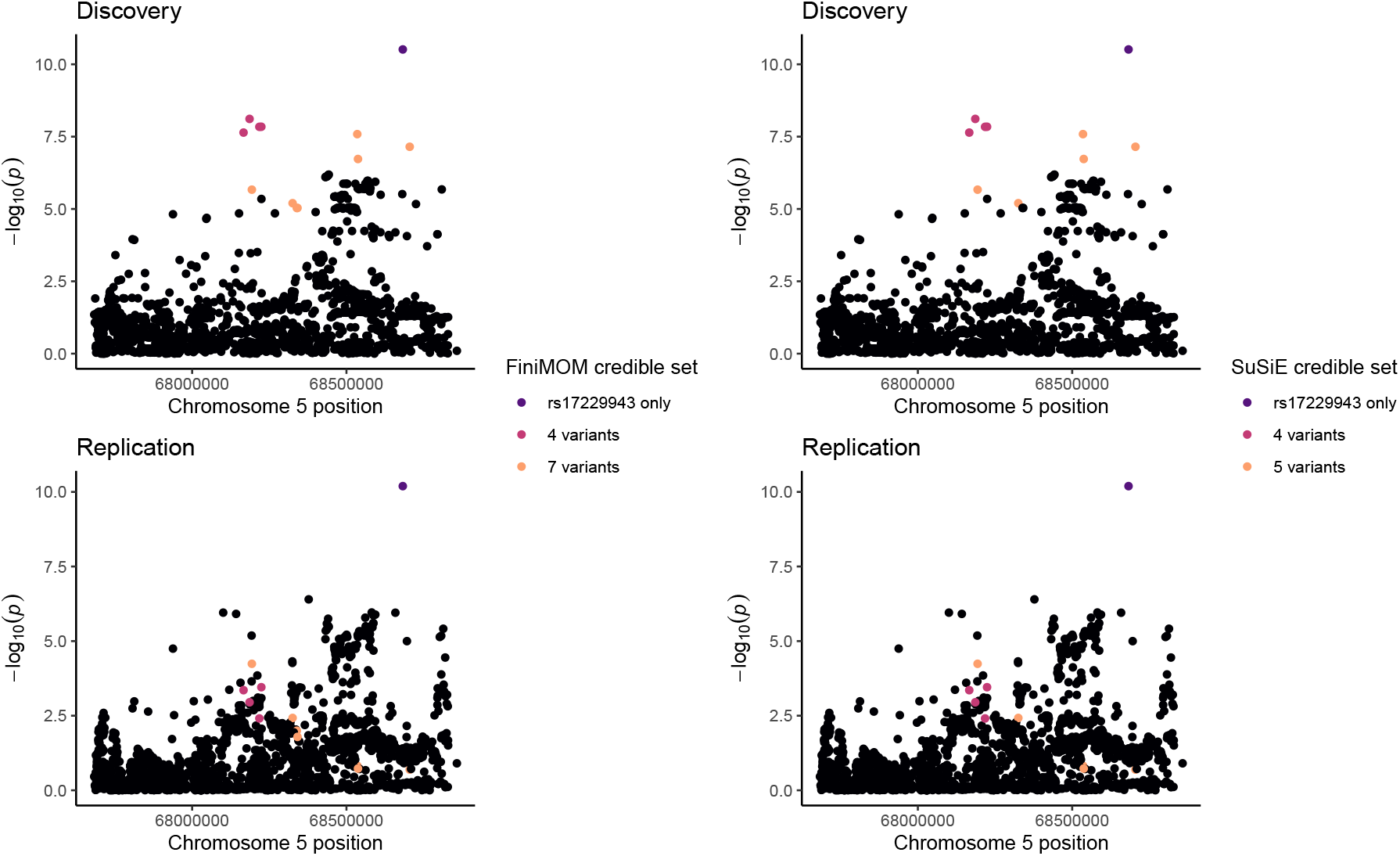
Locus plot of genetic associations (− log_10_(*p*)) per each variant within *RAD17* locus (± 1Mb from rs17229943 variant), with credible sets highlighted for FiniMOM (left panels) and SuSiE (right panels) in the discovery GWAS (top panels) and in the replication GWAS (bottom panels). FiniMOM: fine-mapping using inverse-moment prior; SuSiE: sum of single effects; GWAS: genome-wide association study.

**Figure 9:**
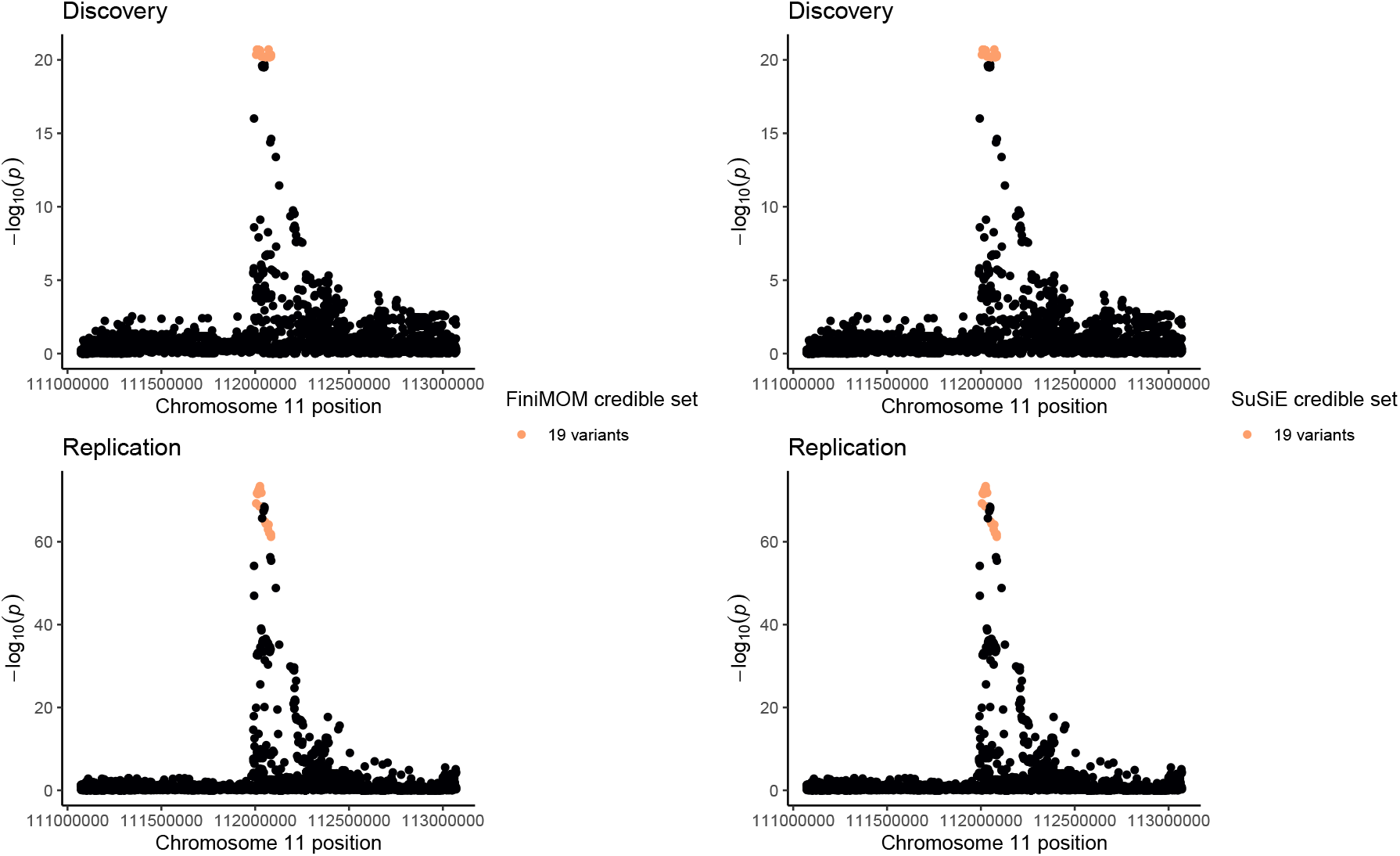
Locus plot of genetic associations (− log_10_(*p*)) per each variant within *BCO2* locus (± 1Mb from rs12420140 variant), with credible sets highlighted for FiniMOM (left panels) and SuSiE (right panels) in the discovery GWAS (top panels) and in the replication GWAS (bottom panels).FiniMOM: fine-mapping using inverse-moment prior; SuSiE: sum of single effects; GWAS: genome-wide association study.

For *NLRC4*, FiniMOM produced a smaller credible set, consisting of only rs385076, the lead variant. This SNP was also the top variant in the replication GWAS. The size of the corresponding SuSiE credible set was 40 variants (Figure 7). Both methods detect variant rs17229943 as the lead signal (and as its own credible set) at *RAD17* locus. In addition, there were two other credible sets detected by both methods. However, no strong signal was detected for the variants in these credible sets in the replication GWAS (Figure 8). Both FiniMOM and SuSiE were able to distinguish the peak *BCO2*, and both produced one credible set with the same variants included. This signal was also detected in the replication GWAS (Figure 9).

## 4 Discussion

In this paper, we have proposed a novel fine-mapping method for summarised genetic associations using product inverse-moment priors. Additionally, incorporating theoretical results of the model dimension prior in very sparse regression settings, we propose an adjustable beta-binomial prior to be used. The proposed approach showed improved detection of causal variants across various scenarios which were aimed to mimic typical settings in pQTL association analyses [44]. Our method also worked well in the applied example, with similar results to SuSiE.

In Bayesian fine-mapping methods, the traditional way is to set a Gaussian prior for the causal effect sizes. This allows for effects that are arbitrarily close to zero with non-negligible probability (Figure 1). While such small effect sizes may exist, these cannot be reliably detected with finite datasets. In contrast, the piMOM prior applied here can be set such that, conditioned on the variant being causal, the probability for the absolute values of the effect sizes being smaller than a specific threshold can be determined *a priori*. Apart from the use of non-local priors [24]–[26], similar approaches can be found in QTL mapping literature [49]–[51], and we believe using such priors is a natural way to analyse genomic datasets.

We also highlight our method’s flexibility in the trade-off between credible set coverage and power, easily incorporated via the hyperparameter *u* that controls the model dimension prior. Similarly, FiniMOM is highly adaptable to deal with either in-sample or out-of-sample LD reference. While the performance of SuSiE does not notably drop while using out-of-sample LD, both coverage and power were weaker compared to FiniMOM in the simulated scenarios. Improved performance of our method over SuSiE was specifically seen in the scenarios of multiple causal variants. Moreover, unlike in SuSiE, but similarly as in FINEMAP method [13], we also obtain the uncertainty in the number of credible sets.

Our novel MCMC sampling strategy is a compromise between full Gibbs and other stochastic methods, such as sure independence screening (SIS) [23], [52]. As a departure to stochastic methods, our method is fully Bayesian in the sense that the full posterior is explored. Therefore, we avoid the problem of different iterative, stochastic, or stepwise models where minor deviations in data may lead to notably different solutions for the final model [53].

Some limitations of our method should be mentioned. Improving the software implementation warrants further work, as the current implementation of FiniMOM is considerably slower than SuSiE, which may become an issue if fine-mapping is intended to be conducted on a large number of phenotypes or loci. In the absence of in-sample LD information, the importance of a good LD reference cannot be overstated and, while our method is shown to be robust for out-of-sample LD references, it naturally cannot recover poor LD reference and data quality. We did not consider the situations where the effect sizes depend on the allele frequencies of the variants [54], however this can be easily incorporated by allowing the prior parameter *τ* to vary across variants [21]. Finally, our current method does not cover varying LD structures across different ancestries [55], infinitesimal polygenic effects [56], or multivariate outcomes [57].

In summary, we have proposed a novel genetic fine-mapping method for summarized data that outperforms a current state-of-the-art fine-mapping method. The specific strengths in detecting multiple causal variants and adaptability to deal with out-of-sample LD information make FiniMOM an attractive option for fine-mapping studies of quantitative traits.

## 6 Acknowledgements

This project has been funded by the Academy of Finland Profi 5 funding for mathematics and AI: data insight for high-dimensional dynamics [Project 326291] (VK, IL, MJS); and European Union’s Horizon 2020 research and innovation programme under Grant Agreement No 848158 [EarlyCause] (VK, SS, MRJ).

We thank all NFBC1966 and NFBC1986 cohort members and researchers who participated in the studies. We also wish to acknowledge the work of the NFBC project center.

## 7 Author contributions

VK, IL and MJS conceptualised the study. VK performed the statistical analysis and drafted the manuscript. All authors interpreted the results, critically revised the manuscript for intellectual content, and approved the submitted version.

## 8 Web resources

Finimom R package: https://vkarhune.github.io/finimom.

SuSiE R package: https://stephenslab.github.io/susieR/.

IL18 GWAS summary statistics: https://doi.org/10.5523/bris.3g3i5smgghp0s2uvm1doflkx9x (discovery); https://doi.org/10.5281/zenodo.2615265 (replication).

## 9 Data availability

NFBC1966 and NFBC1986 genotype data are available by application via http://oulu.fi/nfbc/.

