## Supplementary File for "Genetic fine-mapping from summary data using a non-local prior improves detection of multiple causal variants"

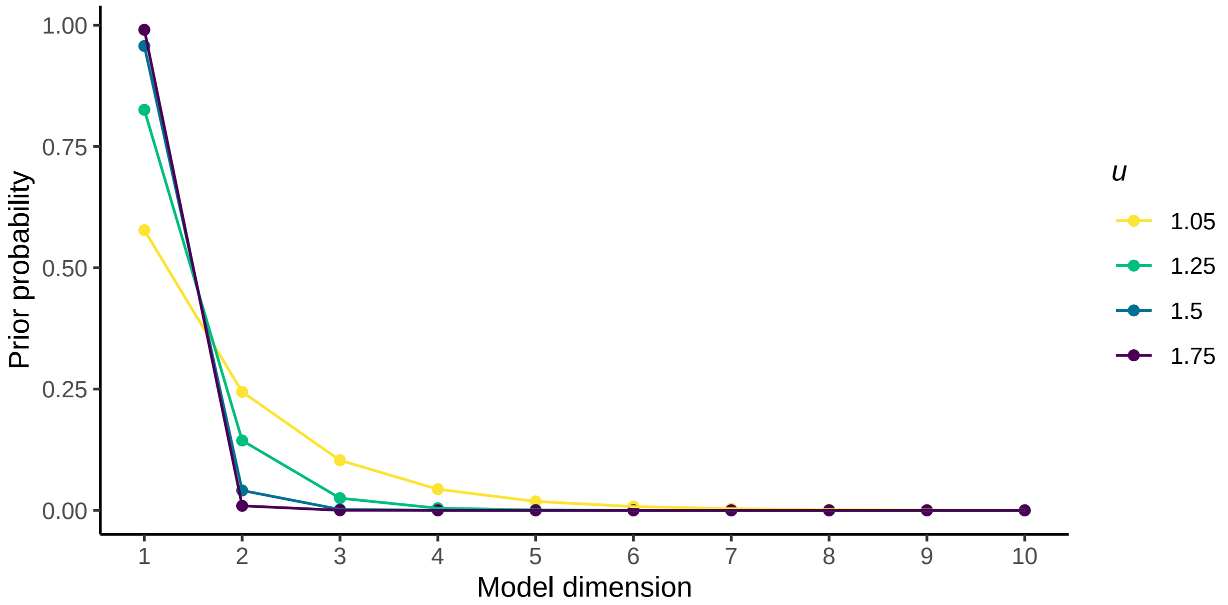


Supplementary Figure 1: Prior distribution for model dimension with different values of $u$, with number of variants $P = 500$ and the maximum model dimension $K = 10$.

Supplementary Table 1: Genomic regions used in the simulations.

| Gene | Chromosome | Start* | End* | Number of variants |
|---|---|---|---|---|
| *UMPS* | 3 | 124349213 | 124564040 | 723 |
| *RPS14* | 5 | 149722753 | 149929319 | 487 |
| *GCNT2* | 6 | 10392456 | 10729601 | 840 |
| *CSNK1A1L* | 13 | 37577398 | 37779803 | 387 |
| *MDGA2* | 14 | 47211134 | 48243999 | 2996 |

* ±100 kb from each gene, given in hg19 coordinates.

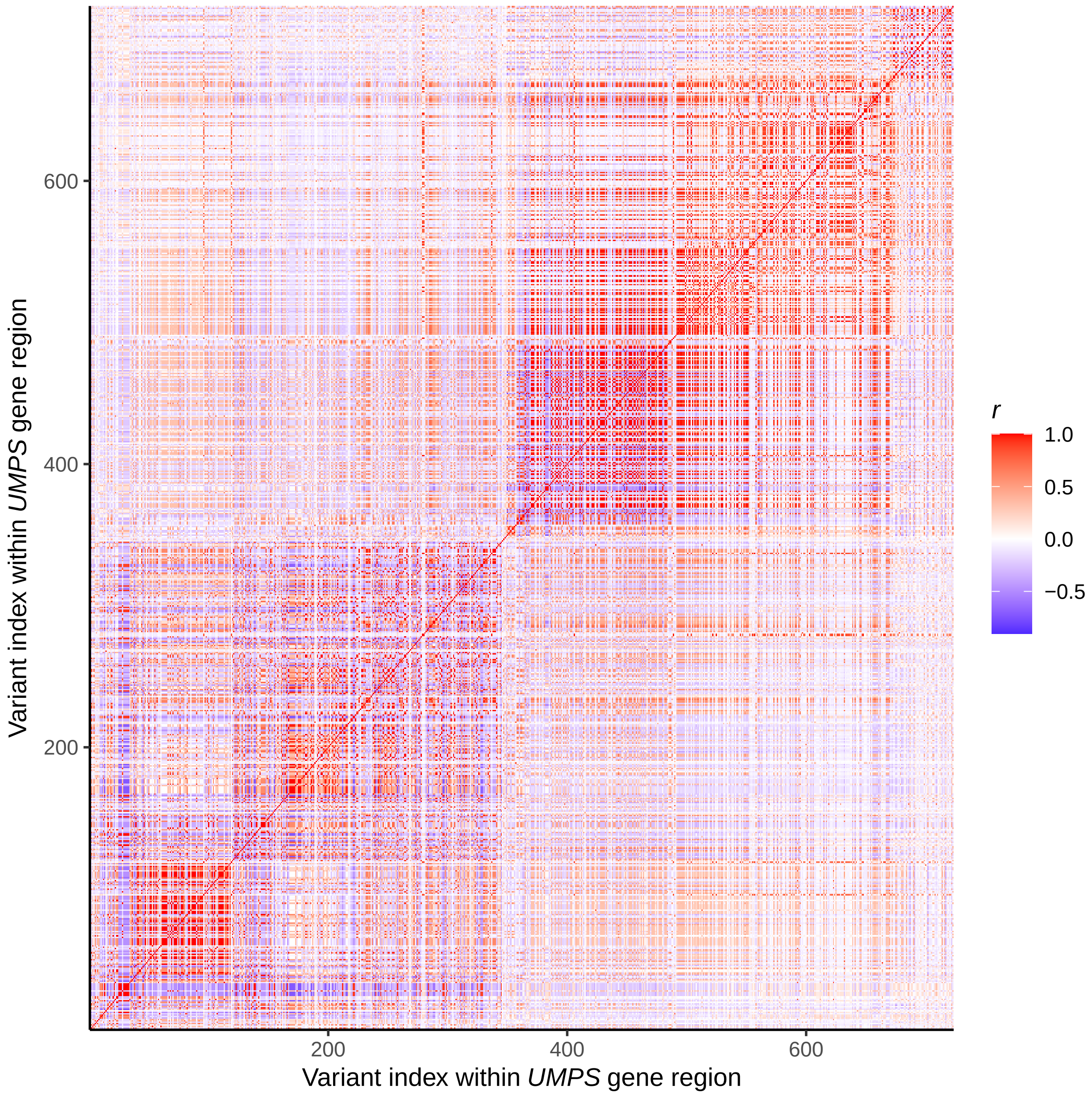


Supplementary Figure 2: Correlation structure within *UMPS* gene region.

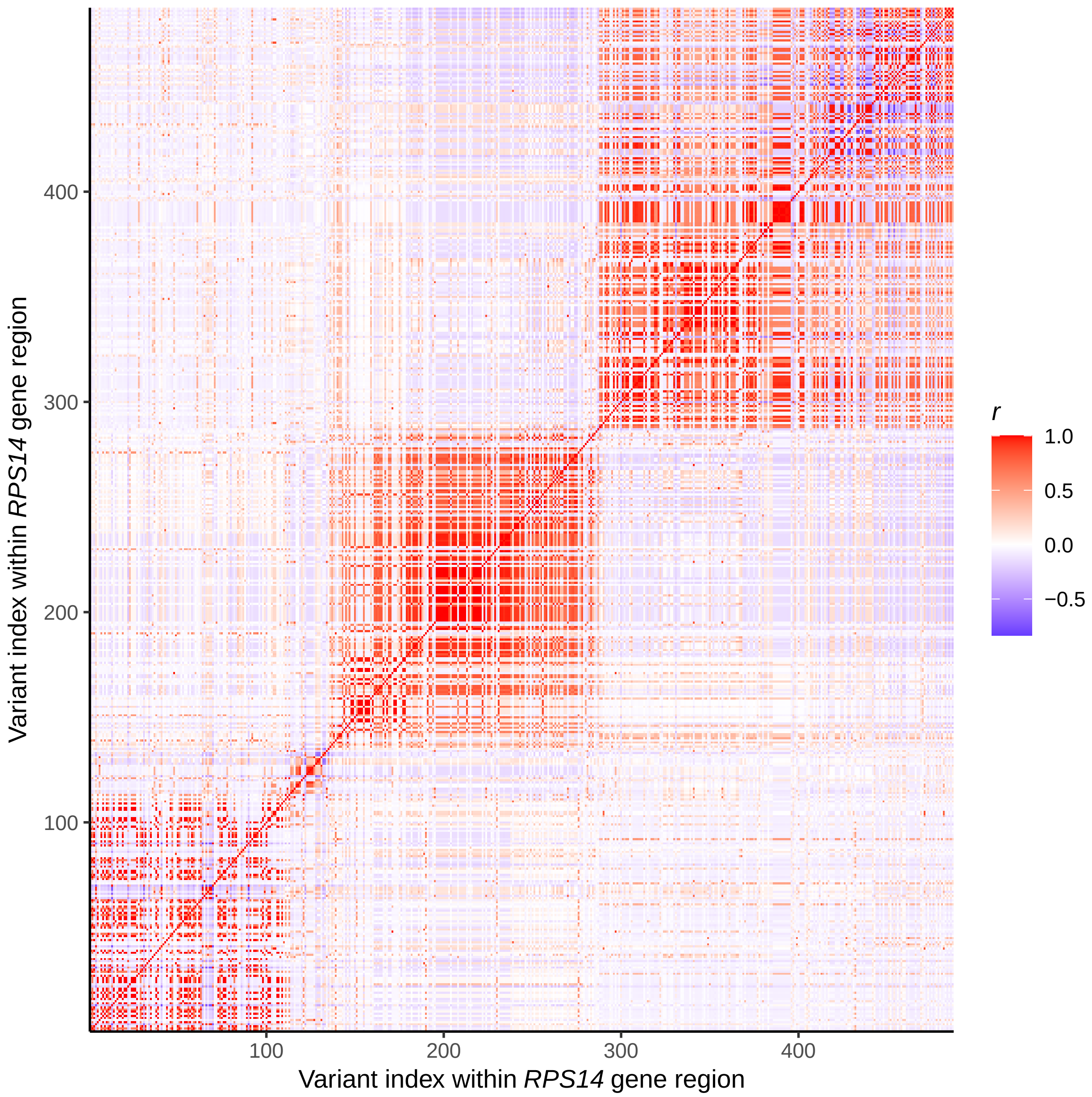


Supplementary Figure 3: Correlation structure within *RPS14* gene region.

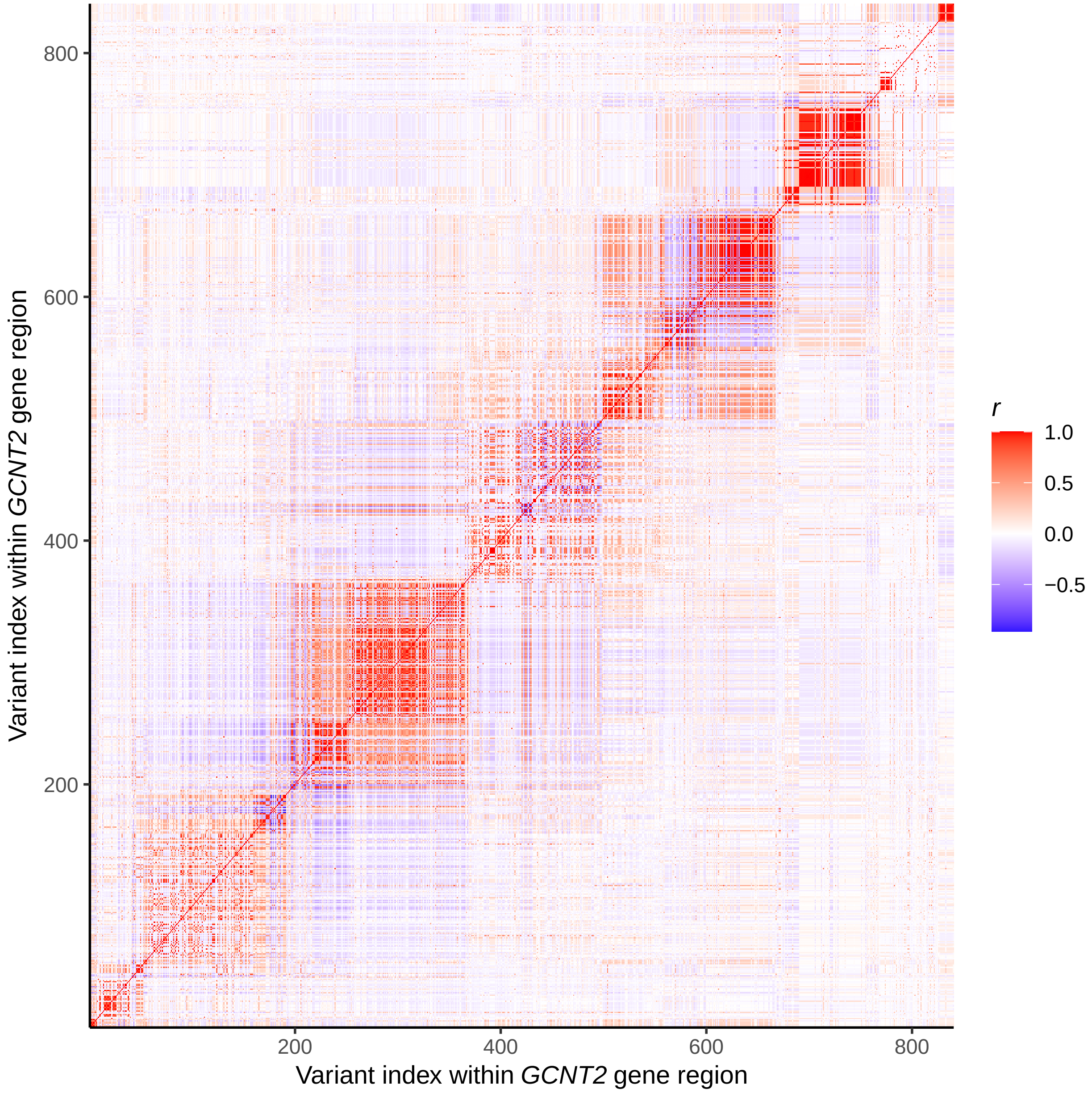


Supplementary Figure 4: Correlation structure within *GCNT2* gene region.

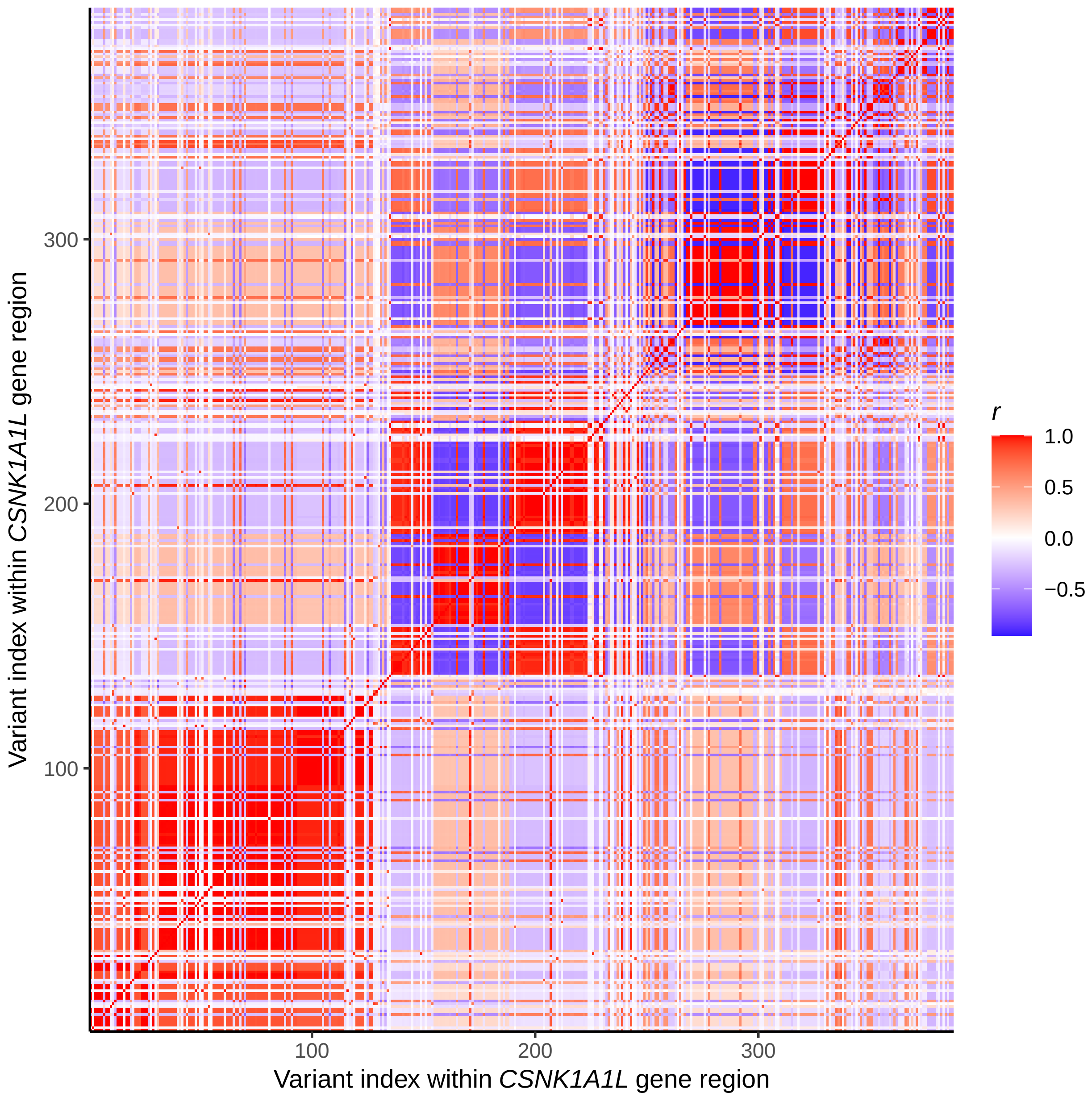


Supplementary Figure 5: Correlation structure within *CSNK1A1L* gene region.

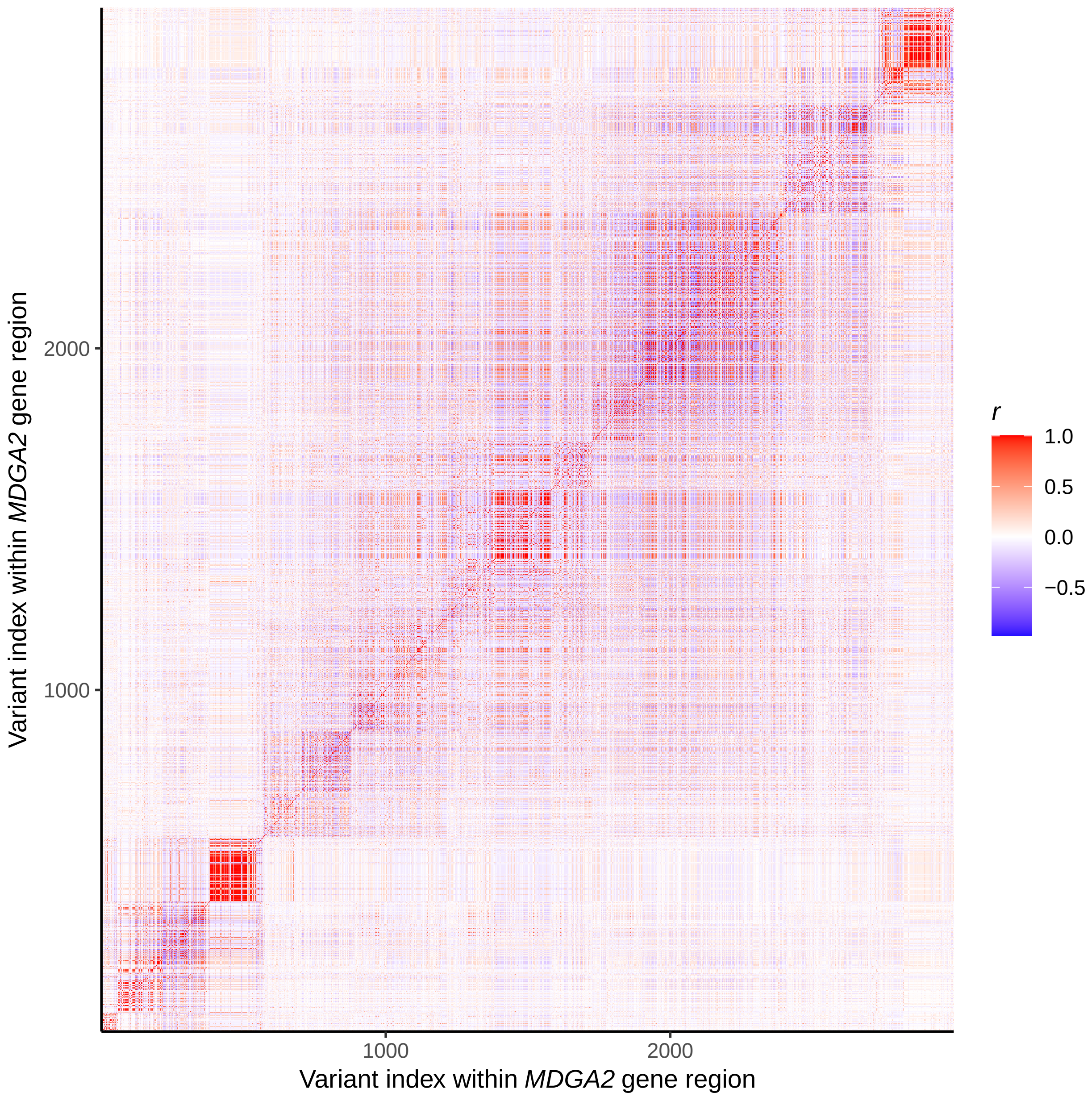


Supplementary Figure 6: Correlation structure within *MDGA2* gene region.
